# Polyploidy impacts population growth and competition with diploids: multigenerational experiments reveal key life history tradeoffs

**DOI:** 10.1101/2022.10.31.514602

**Authors:** Thomas J. Anneberg, Elizabeth M. O’Neill, Tia-Lynn Ashman, Martin M. Turcotte

**Affiliations:** Department of Biological Sciences, University of Pittsburgh, Pittsburgh, PA, USA 15260

**Keywords:** Araceae, duckweed, nutrient limitation, synthetic polyploid, whole genome duplication

## Abstract

- Ecological theory predicts that early generation polyploids (“neopolyploids”) should quickly go extinct owing to the disadvantages of rarity and competition with their diploid progenitors. However, polyploids persist in natural habitats globally. This paradox has been addressed theoretically by recognizing that reproductive assurance of neopolyploids and niche differentiation can promote establishment. Despite this, the direct effects of polyploidy at the population level remain largely untested even though establishment is an intrinsically population-level process.
- We conducted population-level experiments where investment in current and future growth was tracked in four lineage pairs of diploids and synthetic neopolyploids of the aquatic plant *Spirodela polyrhiza*. Population growth was evaluated with and without competition between diploids and neopolyploids across a range of nutrient treatments.
- Although neopolyploid populations produce more biomass, they reach lower population sizes, and have reduced carrying capacities when growing alone or in competition across all nutrient treatments. Thus, contrary to individual-level studies, our population-level data suggest that neopolyploids are competitively inferior to diploids. Conversely, neopolyploid populations have greater investment in dormant propagule production than diploids.
- Our results show that neopolyploid populations should not persist based on current growth dynamics, but high potential future growth may allow polyploids to establish in subsequent growing seasons.

## Introduction

Whole-genome duplication is a common macromutational process across the tree of life (Doyle & Coate, 2019; Fox *et al*., 2020), resulting in the formation of an incipient “neopolyploid” within an otherwise diploid population. At the local scale, neopolyploid individuals are expected to almost always go extinct due to competitive exclusion by their diploid progenitors in their shared niche through mechanisms such as minority cytotype exclusion (Levin, 1975; Husband, 2000; Arrigo & Barker, 2012). However, the expectation of rapid extinction is discordant with the observation that established populations of polyploid plants persist globally in natural environments, with polyploidy acting as a common speciation mechanism in plants (Spoelhof *et al*., 2017; Rice *et al*., 2019). This establishment paradox has been addresssed via theoretical models that focus on population-level dynamics which emphasize the importance of asexuality and niche shifts in promoting establishment success (Rodriguez, 1996; Rausch & Morgan, 2005; Oswald & Nuismer, 2011; Fowler & Levin, 2016; Spoelhof *et al*., 2020b). However, nearly all empirical tests of the ecological effects of polyploidy have been conducted at the individual level. This creates a significant gap in our understanding of polyploid establishment because the performance of an individual does not necessarily equate to the performance of the population. In particular, neopolyploids may differ from diploids in population intrinsic growth rate as well as in their sensitivity to intraspecific competition with themselves or interspecific competition with their diploid ancestors (Hart *et al*., 2018). Moreover, neopolyploidy may alter life-history strategies, such as investment in actively growing progeny versus storage that affects current and future population growth, respectively (Fig. 1a) (Van Noordwijk & Dejong, 1986; Stearns, 1989). The lack of experiments on the immediate effects of neopolyploidy on population-level processes is a crucial missing link to understanding the persistence and establishment of polyploid lineages.

**Fig. 1:**
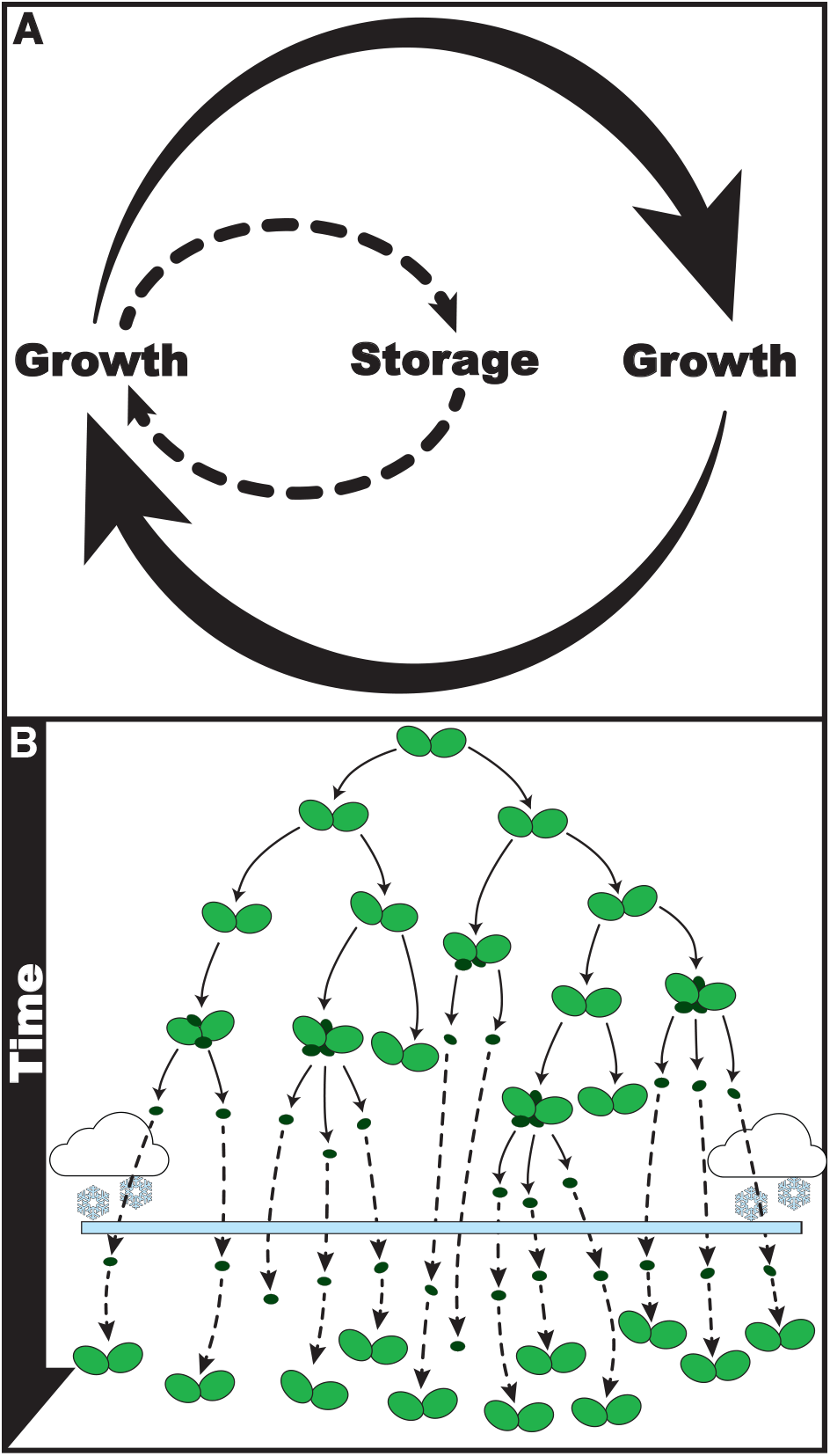
(a) Populations grow by producing active or dormant propagules, both contribute to long term growth. b) The greater duckweed (*Spirodela polyrhiza*) populations propagate via multiple generations (black arrows) of actively growing fronds (green plants) over a growing season and produce turions (black circles) that are dormant propagules that can overwinter (dotted arrows that pass through the blue bar) and regrow to produce future populations of fronds.

At the individual-level, neopolyploidy can lead to instantaneous phenotypic differentiation from their diploid progenitors (Ramsey & Schemske, 2002; Clo & Kolar, 2021). Even when strictly considering autopolyploidy, which does not include the effects of interspecific hybridization, the direct effect of increased genomic content (“nucleotypic” effects) and gene dosage effects (Bomblies, 2020) can lead to immediate phenotypic novelty, such as slower cell division, larger cells (e.g., stomates), or increased body size (Doyle & Coate, 2019; Bomblies, 2020). Neopolyploidy can also lead to phenotypic shifts in the reproductive traits of individuals, such as altered reproductive phenology, increased seed production or clonality (Ramsey & Ramsey, 2014; Van Drunen & Husband, 2018a; Anneberg & Segraves, 2020), which could dramatically affect future population recruitment rates. Additionally, these novel phenotypes can vary with the genotype of the progenitor diploid (Van Drunen & Husband, 2018b; Doyle & Coate, 2019; Wei *et al*., 2020). Thus, the genetic origin of a neopolyploid may be as important as selective tuning and later adaptation in deteriming the success of neopolyploid populations. It is therefore critical to study the effect of neopolyploidy on population performance by using multiple genome duplication events from unique diploid genotypes.

The increased body size associated with neopolyploidy at the individual level (Otto, 2007) is predicted to be adaptive when neopolyploids directly compete with their smaller-bodied diploid ancestors, but there may be population-level costs associated with this individual-level benefit that are rarely considered. For instance, larger-bodied neopolyploids may have a stronger *per capita* competitive effect on their diploid progenitors (Levin, 1983; Hin & de Roos, 2019), but larger sized individuals could also imply stronger intraspecific competition and hence a lower carrying capacity. This would be consistent with individual-level studies showing neopolyploid growth and fitness requires greater nutrient supplies than their diploid progenitors (Guignard *et al*., 2017; Walczyk & Hersch-Green, 2019; Anneberg & Segraves, 2020). Given that neopolyploids often have higher nutrient requirements, we expect them to have slower population growth and should be more sensitive to diploid competition when resources are limited as a result (Guignard *et al*., 2017; Hart *et al*., 2018). These patterns could lead to a trade-off between increased biomass production and population abundance. For instance, metabolic scaling theory predicts that while larger individuals have relatively lower metabolisms and grow more slowly than small individuals, they have one demographic advantage - their metabolic demands per unit mass are lower so populations can achieve higher biomass (Marshall, 2022). Additionally, the ratio of nutrient supply (e.g., nitrogen (N): phosphorus (P)) and the relative differences in requirements for these nutrients by diploids and neopolyploids may alter competitive outcomes (MacArthur, 1972; Tilman, 1982). Thus, by manipulating not only the concentration, but also the ratio of nutrients, we can test whether nutrient stoichiometry can mediate the competitive dynamics between neopolyploids and their diploid ancestors. This may be especially important if whole-genome duplication causes a niche shift, reducing competition among the ploidal types and the likelihood that neopolyploids go extinct in all resource environments (Rodriguez, 1996; Oswald & Nuismer, 2011; Fowler & Levin, 2016).

While previous individual-level studies focus on current productivity (e.g., biomass) (Collins *et al*., 2011; Thompson *et al*., 2015), none to our knowledge have explored how alternative life-history strategies are affected by competition. Beyond growth in the current season, many species can facultatively engage in storage by banking dormant individuals for future growth, especially when resources become scarce (Venable & Brown, 1988). Examples include egg or seed banking, investment into rhizomes, and recalcitrant spore production (Baskin & Baskin, 2014; Martinez-Garcia & Tarnita, 2017). At the individual level, neopolyploidy can lead to an immediate increase in storage investment (e.g., root buds or seed mass) (Van Drunen & Husband, 2018a; Anneberg & Segraves, 2020). As such, neopolyploid persistence may be mediated by both the timing and total production of stored propagules, given that they could circumvent times of low nutrient supply or competitively stressful growing conditions through dormancy (Caceres, 1997; Angert *et al*., 2009). Furthermore, whether the competitive interactions between diploid and neopolyploid populations in current (active) growth mirror those on future growth potential (dormant propagules) remains untested.

To fill these gaps in knowledge, we ask 1) Is there an advantage to neopolyploidy at the population level, and does it depend on resource availability or the genetic background of the progenitor diploid? 2) When growing together, do neopolyploid or progenitor diploid populations have a competitive advantage over the other in growth and abundance? 3) Do the determinants of ploidal-specific investment into future population growth mirror current population growth? We addressed these questions by synthesizing neopolyploids from four genetically distinct diploid genotypes of a floating aquatic plant, the ‘greater duckweed’ *Spirodela polyrhiza* (Araceae; SI Extended Methods) that reproduces asexually through either clonal division of actively growing fronds or dormant propagules (turions) (Appenroth & Nickel, 2010) (Fig. 1b). We chose to study an asexually reproducing species since current theory predicts that reproductive assurance, such as via clonal reproduction, can alleviate minority cytotype challenges and contribute to neopolyploid establishment (Fowler & Levin, 2016; Van Drunen & Husband, 2019; Spoelhof *et al*., 2020a). The potential importance of asexual reproduction in polyploid establishment is also supported in empirical reviews (Van Drunen & Husband 2019). As a result, *S. polyrhiza* is a biologically relevant and tractable system to address the factors governing polyploid establishment especially at the population level. Additionally, we used *S. polyrhiza* since it is one of the fastest reproducing macroscopic plants (Ziegler *et al*., 2015), which allows us to conduct large replicated multigenerational population dynamic experiments. The asexual reproductive strategy of *S. polyrhiza* also allows us to avoid the confounding effects of other mechanisms such as outbreeding between diploids and neopolyploids that can contribute to minority cytotype exclusion (Levin, 1975) and impact respective population dynamics. In this study, we evaluated replicated multigenerational population dynamics of neopolyploids and their progenitor diploids when grown both alone or in competition across a range of N and P that varied in concentration and stoichiometry. Synthetic neopolyploids are recognized as a powerful tool for studying the direct effects of genome duplication because they avoid the confounding effects of evolution after duplication that exist in wild polyploid-diploid comparisons (Bomblies, 2020). Furthermore, including multiple genotypes of synthetic neopolyploids allows us to discern how repeatable the immediate effects of genome doubling are when considering the standing intraspecific genetic diversity of progenitor diploids.

## Materials and Methods

See Methods S1 for an extended description of all methods.

### Study system and synthesis of neopolyploids

*Spirodela polyrhiza* (Araceae) is a model system for testing multigenerational population dynamics across ecological settings (Lam *et al*., 2014; Wang *et al*., 2014; Laird & Barks, 2018; Hart *et al*., 2019). This freshwater aquatic plant has a relatively small genome size (158 Mbp) with rapid generation times (Ziegler *et al*., 2015) that almost always reproduce clonally and very rarely flower (Jacobs, 1947). Additionally, their fast generation time to maturity (2-3 days) allows us to track multigenerational population dynamics within a couple of weeks. *S. polyrhiza* reproduce clonally via: 1) the production of actively growing fronds that separate from their maternal plant and contribute to current-season populations, and 2) production of dormant propagules (turions) that overwinter and contribute to future populations (Jacobs, 1947; Appenroth & Nickel, 2010) (Fig. 1b). Since turions are produced in response to stress (e.g., nutrient scarcity or onset of winter) and remain dormant until growing conditions become favorable (i.e., transition from winter to spring), their production can be considered investment in future growth (Appenroth & Nickel, 2010).

Four genetically distinct ancestral diploid lineages of *S. polyrhiza* were collected in western Pennsylvania, USA (Table S1 for collection site info). These diploid lineages are based on several microsatellite markers (Hart et al. 2019; Kerstetter et al. *In Review)*. In 2019 and 2020 we generated four synthetic neotetraploid (neopolyploid) lineages from each of the four diploid *S. polyrhiza* lineages via application of the mitotic inhibitor colchicine (Sigma Aldrich, CAS: 64-86-8) followed by confirmation of their ploidy level with flow cytometry as outlined in Wei et al. 2020. Population growth rates of colchicine-treated diploids that did not convert to tetraploids were not different from diploids belonging to the same lineage that were only exposed to the treatment solvent (Fig. S1). Nevertheless, to be conservative, the following comparative analyses were conducted on a colchicine-treated but unconverted diploid population versus the corresponding neotetraploid from each lineage.

### Experimental setup

Experimental populations were seeded with six fronds in 950 ml plastic containers filled with 500 ml of modified Appenroth media (Appenroth *et al*., 1996), according to their prescribed nutrient treatment (Table S2 for recipes). Populations grew in a greenhouse in the summer of 2021 at the University of Pittsburgh for 17 days. Since duckweeds have very rapid generation time (2 – 3 days from bud to reproductive maturity), up to 8 generations can occur in 17 days, making this duration on par with other multigenerational population dynamic studies with duckweeds (Armitage & Jones, 2019; Hart *et al*., 2019).

We conducted two experiments. The first experiment tested the monoculture response of three population level traits (timeseries of frond abundance, biomass productivity, and timeseries of turion abundance) to variation in ploidy, genotypic lineage, and N & P concentration. The second experiment analyzed these same parameters with the added contrast of diploids and their neopolyploid descendants being grown either alone or together in an additive design. In both experiments, we counted the total number of actively growing fronds and turions from photographs every two days and measured dry biomass at the final day of the experiments as an estimate of productivity. We grew five replicates of each ploidy level of each lineage and each treatment level in the first experiment, and 10 replicates per treatment level in the second experiment.

### Statistical analyses

We analyzed each of the three population level traits with either linear or generalized linear mixed effect models. In these models, to account for any pre-treatment differences, we included a covariate of initial surface area covered by the six starting individual fronds in each sample. To test the population-level performance in the first experiment, we fit linear mixed models to the frond and turion count data by including a random slope of the crossed effect of genetic lineage by day of experiment. We chose to analyze the frond and turion growth data separately since the dormant turions do not contribute to population growth in the current season. In these models, we specified ploidy, N concentration, P concentration, lineage, and their interactions as fixed effects. We further estimated the carrying capacity for both ploidy levels of each of the four genetic lineages by fitting a logistic growth model to the data with the nls_table function from the forestmangr package (Braga *et al*., 2020) in R (R Core Team, 2021). After log_10_-transformation to improve the condition of normality, we fit the estimated carrying capacity data to a linear model that included ploidy, nutrient treatment, and genetic lineage as factors. We further analyzed differences in productivity between diploid and neopolyploid populations with their respective final dry biomass from day of harvest. Since final dry biomass data measurements had uneven residual errors, we analyzed these data with a generalized mixed effect model with a gamma distribution (Bates *et al*., 2015). In all statistical models, we calculated a *post-hoc* least square mean estimate for each treatment level using the lsmeans package (Lenth, 2016) in R (R Core Team, 2021). From the least square means estimates, we derived the relative percent increase or decrease of neopolyploid population performance in relation to diploids.

We analyzed how competition between diploids and neopolyploids affects their population performance through similar timeseries linear mixed models as described above. We fit either linear or generalized mixed effect models that had ploidy, nutrient treatment, lineage, day of experiment, competition, and their interaction effects as fixed effects with the pre-treatment frond surface area as a covariate and a random effect of day of experiment crossed with each genotype of each ploidy level to account for repeated measures. We additionally estimated how the carrying capacity of diploids and neopolyploids of each genetic lineage was affected by competition the same as described above. When estimating carrying capacities from the competition experiment, we observed that populations in high nutrient treatments were still growing too quickly, causing spurious carrying capacity estimates. Thus, we omitted the high nutrient treatment from our main analysis of carrying capacity (all estimates are reported in Appendix 1).

## Results

### Effect of polyploidy on current population growth

The current growth of neopolyploid populations was slower than their diploid progenitors, ultimately reaching smaller population sizes by the end of the experiment across all nutrient treatment levels (Fig. 2; Table S3). For example, averaging over all genetic lineages, neopolyploid population sizes were 42% and 28% lower than their diploids progenitors in the lowest and highest nutrient treatments, respectively. Our estimation of carrying capacity corroborated the current growth data, with neopolyploids having significantly lower estimated carrying capacities than their diploid progenitors across all nutrient treatments (Fig. 3; Table S4). In addition to polyploidy depressing carrying capacity, the significant ploidy-by-day interaction indicates that polyploids are also slower growing in general (Table S3).

**Fig. 2:**
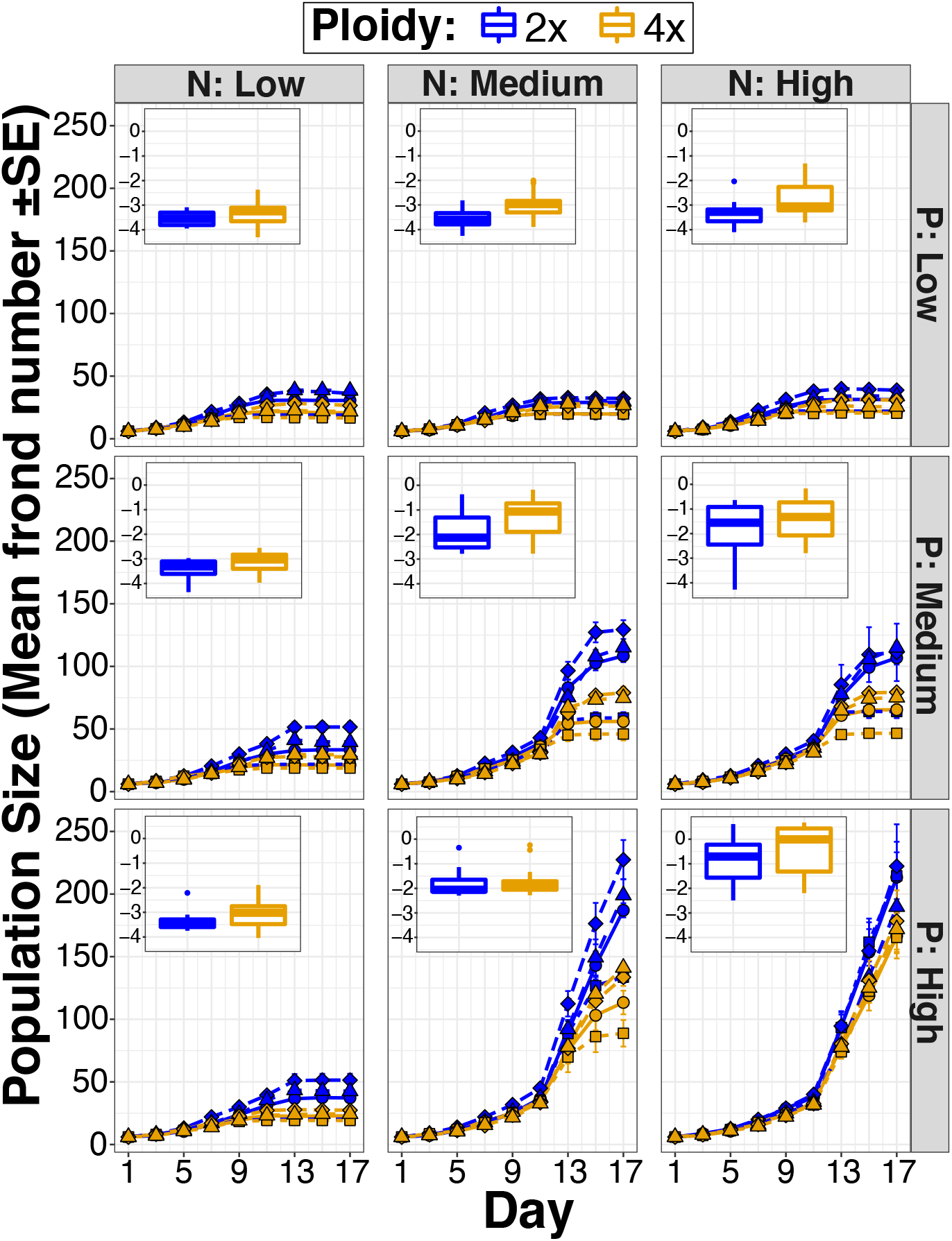
Population sizes (based on counts of number of fronds) of diploids (2x) and neopolyploids (4×) growing separately, across a gradient of nitrogen ‘N’ and phosphorus ‘P’ concentrations for each of the four genetic lineages separately (line type). Neopolyploid populations grow slower and reach smaller population sizes across all nutrient levels and genetic lineages. Boxplot insets show the median and interquartile range of neopolyploid population productivity (log_e_ dry frond biomass) is greater than diploids by day 17 across all nutrient treatments.

**Fig. 3:**
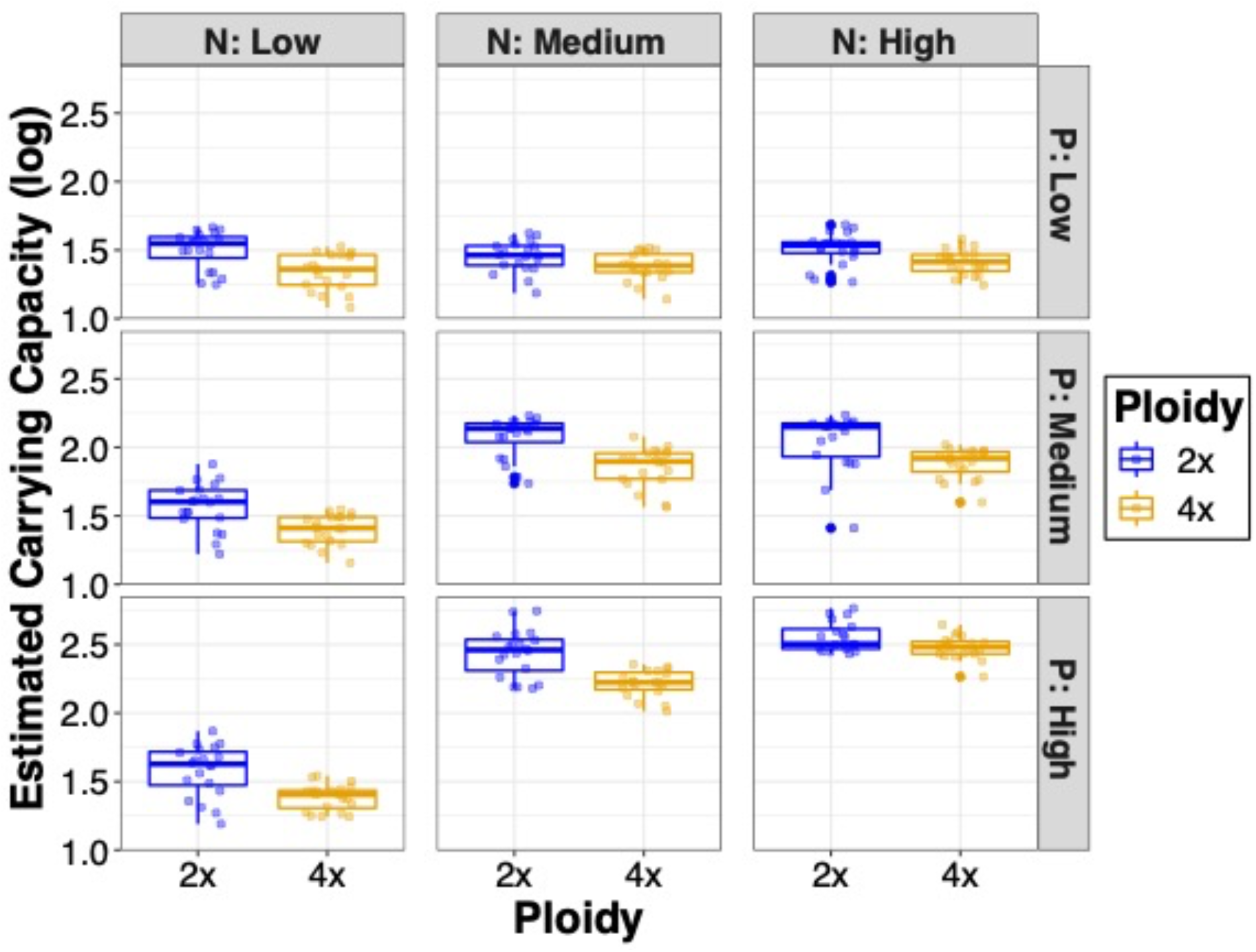
Estimated carrying capacity (boxplots show median and interquartile range) of current growth fronds (log_10_) for diploid (2×) and neopolyploid (4×) populations grown alone across the nine nutrient treatments (Nitrogen and Phosphorus). Replicate populations used to estimate carrying capacity are denoted by the translucent dots overlaid on boxplots.

Despite neopolyploids never reaching larger populations sizes than diploids, neopolyploids responded more to N enrichment: averaging across phosphorus concentrations, neopolyploid population sizes increased 43% more than diploids in response to nitrogen enrichment, but the strength of this pattern was strongly dependent on genetic background (i.e., ploidy, nitrogen, and lineage interaction; *F*_6, 3035_ = 3.00, *P* = 0.006; Table S1). Conversely, diploid population sizes increased by 15% more than neopolyploids in response to P enrichment (i.e., ploidy and phosphorus interaction; *F*_2, 3035_ = 3.27, *P* = 0.038), a pattern that did not vary among the four genetic lineages (Table S3). Yet, diploids and neopolyploids did not differ in their response to variation in the N:P stoichiometric ratio (Table S5). The negative effect of neopolyploidy on population size when grown alone, regardless of nutrient supply or genetic lineage, indicates that the disadvantage of slower population growth is a universal cost of genome duplication for this aquatic plant.

In contrast to population size, and in agreement with predictions from metabolic scaling theory (Marshall, 2022) as well as many individual-level studies (Clo & Kolar, 2021), neopolyploid populations were more productive (final dry biomass) across all nutrient conditions. A least-square mean comparison showed that neopolyploid populations accumulated 85 % more biomass via current growth than diploids (Fig. 2; χ^2^ = 6.23, df = 1, *P* = 0.013; Table S6). Although N and P enrichment significantly increased biomass productivity, polyploidy did not interact with nutrient treatments or genetic lineage on productivity (Fig. 2; Table S6; Fig. S2). Thus, the main effect of polyploidy was a substantial increase in population biomass productivity.

### Competitive dynamics of diploids and neopolyploids

When we grew diploids and neopolyploids either alone or in competition, we showed that diploids grew faster and produced greater population sizes than their neopolyploid descendants, and this pattern held true for each nutrient treatment level and for all four genetic lineages (Fig. 4, Table S7). Our analysis of the estimated carrying capacities corroborated the analysis of population sizes. Neopolyploidy reduced carrying capacity across all nutrient treatments and regardless of competition (Fig. 5; Table S8). Surprisingly, competition between diploids and neopolyploids did not interact with nutrient treatment over the course of the experiment (Table S7). However, the effect of competition between diploids and neopolyploids did strongly interact with genetic background (Fig. 4; *F*_3, 3224_ = 2.877, *P* = 0.035; Table S7), suggesting that the effect of polyploidy on competition is more complex than simple genome size doubling alone.

**Fig. 4:**
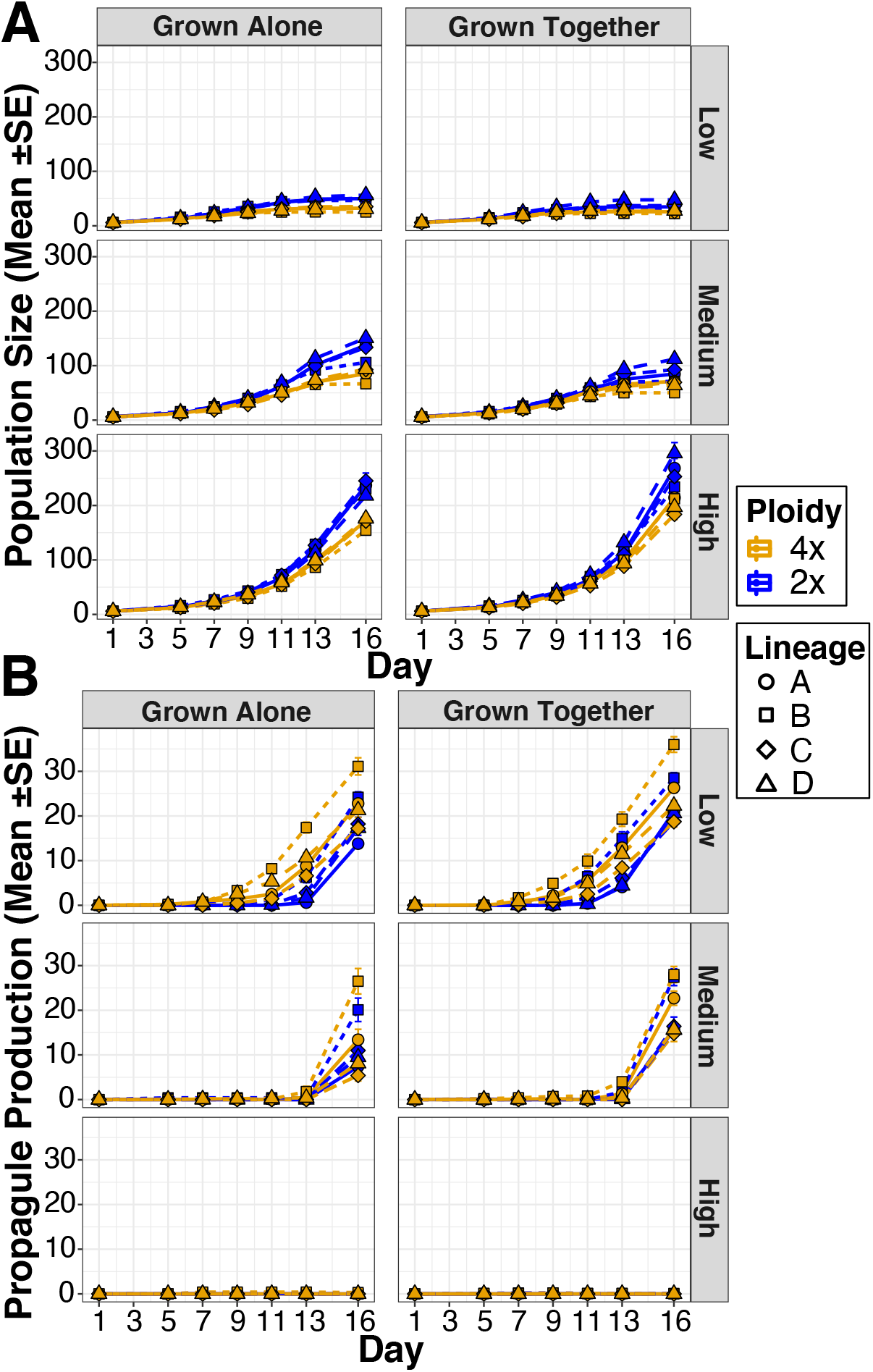
Diploid and neopolyploid population dynamics (frond population size or turion production over day in experiment) when grown alone or together in three nutrient levels for the four genetic lineages (shapes). a) Current population size of diploids (2×) is greater than polyploids (4×) in all treatments, regardless of competition. b) Neopolyploid populations had earlier and greater investment in turions than diploids, and competition caused both ploidy levels to increase propagule production.

**Fig. 5:**
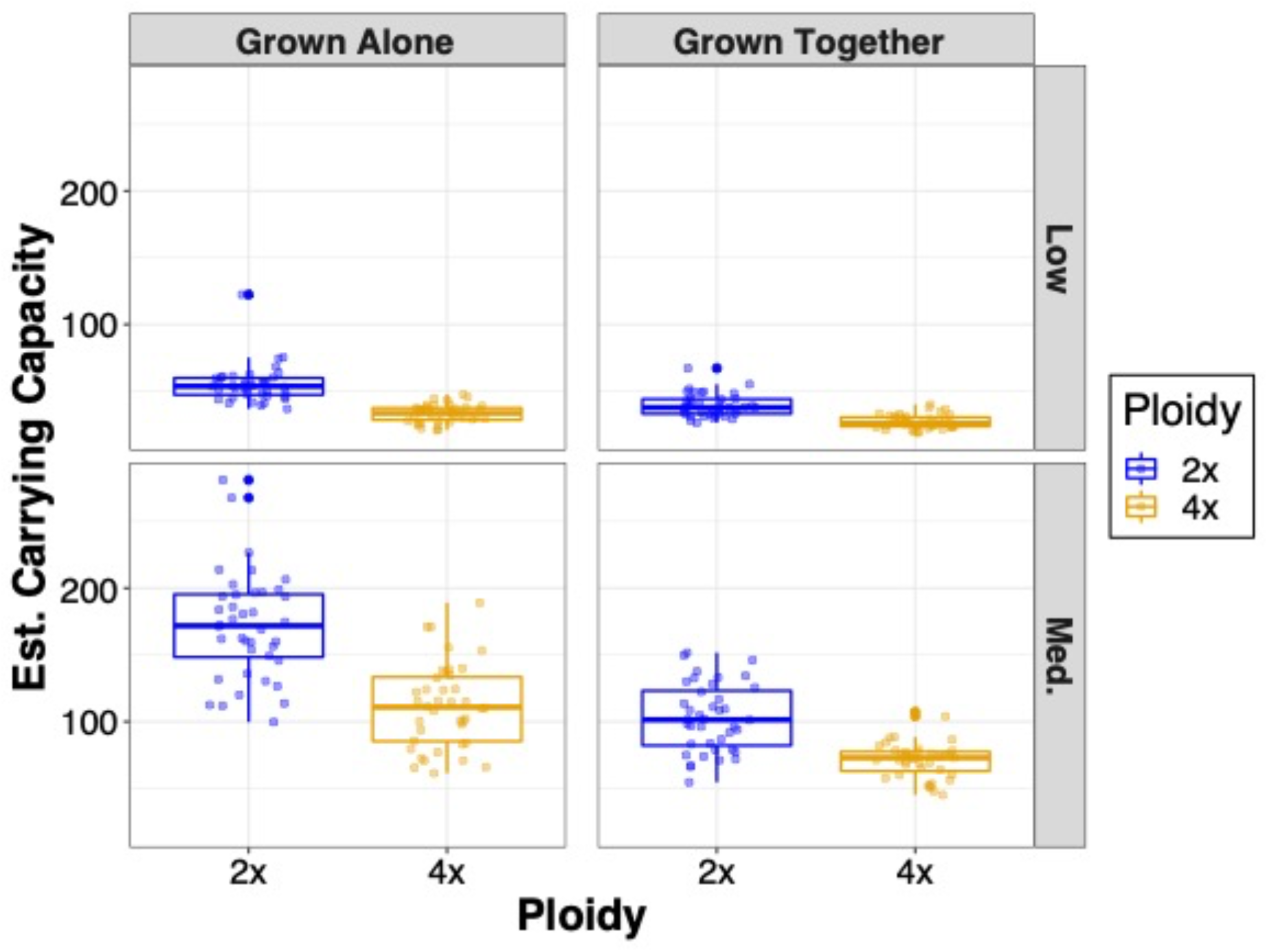
Estimated carrying capacities of current growth fronds of diploid and neopolyploid populations grown in monoculture or competition and either in low or medium nutrient treatment. Individual populations from which we estimated carrying capacity are denoted by the translucent dots overlaid on boxplots. High nutrient treatment samples not shown due to poor fit to a logistic growth model.

While neopolyploid populations again produced more biomass than diploids in monoculture (37 % greater final dry biomass than diploids; Fig. S3), competition caused the biomass productivity of neopolyploids to decline more than diploids. However, neopolyploids still produced more biomass in competition than diploids overall, especially with high nutrient supply (neopolyploid biomass productivity decline of 65 % in high nutrients versus 47 % in low nutrients; Fig. S3; Table S9; χ^2^ = 13.42, df = 2, *P* = 0.001; Table S10), and this pattern was consistent among genetic lineages (χ^2^ = 7.11, df = 6, *P* = 0.31; Table S10).

### Ploidal-specific investment into future population growth

Unlike current (frond) growth, neopolyploids outperformed diploids in future (turion) growth potential when grown as monocultures (Figure 1B). Neopolyploid populations produced turions earlier and accumulated more of these dormant propagules than diploids, especially in lower nitrogen environments (Fig. 6; Table S11, S12). For instance, when grown alone across the nine nutrient levels, neopolyploid populations started producing turions on average six days earlier in response to low nitrogen supply (χ^2^ = 6.01, df = 2, *P* = 0.05; Table S11). A separate generalized model of the total turion production over time found that the effect of neopolyploidy was highly dependent on N and P concentrations, in which neopolyploids produced significantly more turions in lower nutrient environments (χ^2^ = 21.59, df = 4, *P* ≤ 0.001; Table S12). Furthermore, there was no interaction between ploidy level and genetic lineage on turion production over the course of the experiment (*F*_3,49_ = 1.457, *P* = 0.239).

**Fig. 6:**
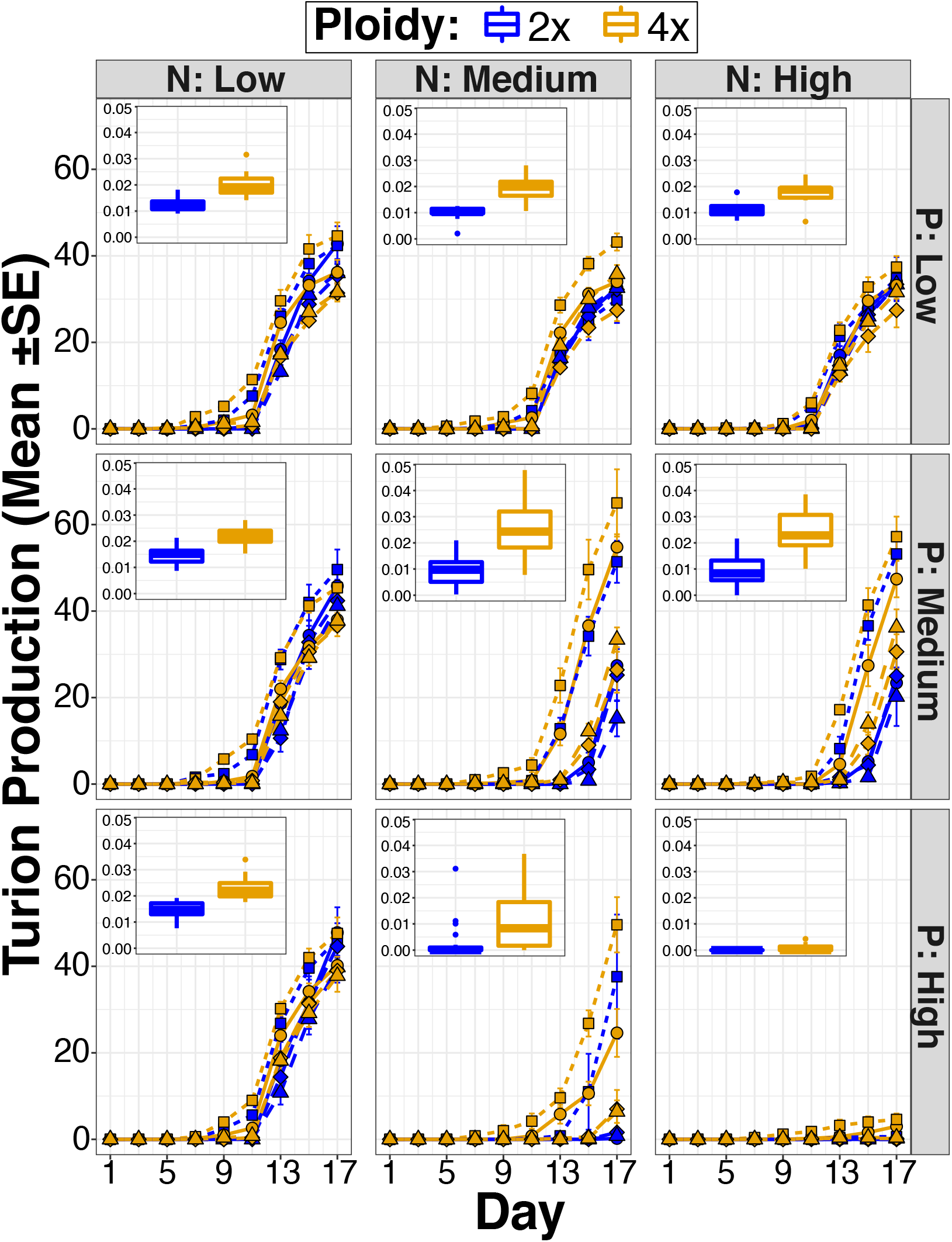
Future growth investment (in number of turions produced) of diploid (2x) and neopolyploid (4x) populations when grown alone. Insets show the loge grams dry biomass of turions across various N and P concentrations. Neopolyploid populations produced more turions than diploids, and this pattern depends on genetic lineage (shapes as in Fig. 3).

When grown with or without competition, neopolyploid populations again produced more turions and began production earlier (Fig. 4). Interestingly, competition elicited a disproportionately greater investment in storage by neopolyploid populations under medium nutrient supply than diploids, whereas diploid populations increased turion production disproportionately more with low nutrients (χ^2^ = 49.33, df = 2, *P* < 0.001; Fig. 4; Table S13). Nutrient supplies strongly dictated which ploidy level produced more turions in response to competition. A least-square means comparison of turions produced from populations under competition relative to monoculture showed that diploid populations increased turion production more than neopolyploids under low resources (diploid increase 25 % versus 12 % for neopolyploids), but neopolyploids increased turion production more than diploids in medium nutrient environments (neopolyploid increase of 75 % versus 67 % for diploids).

## Discussion

Our population-level experiments on growth and competitive dynamics revealed not only a hidden cost – larger bodied neopolyploids grow more slowly and reach lower carrying capacities overall (Fig. 2; Fig. 3; Fig. 4; Fig. 5), but also a hidden advantage — higher population productivity — of polyploidy. We also revealed an important change in life history strategy as neopolyploids shift toward an increased investment in future population growth potential (Fig 4 & 6). This allocation shift was evident not only at the population level but also at the individual level, since neopolyploid fronds produced more turions *per capita* than their diploid ancestors (Fig S4). Furthermore, neopolyploid populations began to produce more dormant propagules much earlier than their diploid progenitors, a strategy that may provide polyploid populations the ability to readily establish after seasonal deterioration (i.e., winter) or unexpected harsh conditions. Recent theoretical work has predicted that establishment success of neopolyploid populations is strongly dependent on the number of propagules they can produce when growing in competition with their diploid progenitors (Fowler & Levin, 2016; Levin, 2021). We provide evidence to support this expectation and importantly that the population-level patterns we detected vary with their genetic background and thus are not simply a product of genome doubling *per se*, but may also be the result of a higher-level genetic effect of neopolyploidy (Osborn *et al*., 2003).

If establishment and persistence of neopolyploids depends only on current population growth and carrying capacities (Fig 1A), then our results indicate they would likely be eliminated over time across the broad range of resource environments tested. This was indicated by not only reduced population growth rates of neopolyploids (Fig. 2), but also lower carrying capacities (Fig. 3) over the duration of six – eight generations. In fact, future modeling of the conditions for polyploid establishment should incorporate unequal carrying capacities rather than the often-assumed equality. Surprisingly, although we expected competition to favor populations of neopolyploids over their diploid progenitors, we found that the current growth of neopolyploid populations was inferior to diploids, regardless of competition (Fig. 4). However, even though the population sizes of both neopolyploids and diploids decreased in response to competition in lower nutrient treatments, they both grew in population size and maintained positive carrying capacities. Although we did not observe competitive exclusion, to thoroughly test whether diploids and polyploids can coexist would require addition experiments specifically designed to test coexistence (reviewed in Godwin et al 2020), as conducted by Hart et al. (2019) with *S. polyrhiza* competing against another duckweed species.

Counter to our current growth data on population sizes, neopolyploids were more productive in terms of accumulated biomass than their diploid ancestors (Fig. 2 insets). Since neopolyploid populations consisted of larger-bodied fronds that were more productive regardless of nutrient treatment (Fig. 2), our results are consistent with metabolic size scaling rules for larger cells (Marshall, 2022) in which larger cells are metabolically more active but also more efficient. Although we expected that by providing higher nutrient supplies to neopolyploid populations it would alleviate any growth disadvantage compared to their diploid progenitors, we found that was not the case (Fig. 2). Therefore, although increasing nutrients may benefit neopolyploid individuals (Walczyk & Hersch-Green, 2019; Anneberg & Segraves, 2020), this does not necessarily manifest at the population level. The pattern of neopolyploid populations comprising larger-bodied fronds that achieve lower population sizes overall, highlights a limitation in the individual-based paradigm which often focuses exclusively on biomass productivity when assessing neopolyploid performance. If establishment was based on biomass productivity patterns alone, we would conclude neopolyploids are unambiguously the winner against their diploid progenitors; however, the larger population sizes of diploids reveal that neopolyploids are poorer off and may not persist.

Yet, our results reveal an unexpected mechanism that could provide an early numerical advantage to neopolyploid populations by shifting their life-history strategy (Stearns, 1989), which could allow establishment. We found that neopolyploid populations produced more dormant propagules (turions) and began earlier than diploids, investing more heavily into their future population growth. This is an under-appreciated mechanism that could help explain how polyploids establish and why mixed-ploidy populations of species are common in nature (Duchoslav *et al*., 2010; Kolar *et al*., 2017) as storage of dormant individuals can be a mechanism of coexistence among competing taxa (Warner & Chesson, 1985; Caceres, 1997; Angert *et al*., 2009; Armitage & Jones, 2019). Specifically, environments with oscillations between benign and harsh growing conditions may lead to a situation where polyploid establishment is favored since neopolyploids invested more in future growth than diploids. If neopolyploid populations overwinter with more dormant propagules than diploids, they could have a numerical advantage in the spring which could allow either coexistence or exclusion of their diploid progenitors. Additionally, the generally greater investment into future growth and earlier initiation of it by neopolyploids constitutes a shift towards a more conservative risk-averse strategy that would allow them to circumvent hostile growing periods such as an early onset freeze (Jacobs, 1947; Childs *et al*., 2010). This shift could be one explanation of the global pattern of established polyploids in polar and stressful habitats and with their evolutionary resilience to cataclysmic climate events, such as the K-T extinction (Madlung, 2013; Van De Peer *et al*., 2017) and the relative absence of established polyploids in the tropics with a low degree of seasonality (Rice *et al*., 2019). Our future growth data, showing that neopolyploids produce larger and more dormant propagules than their diploid ancestors is an important step in understanding the factors promoting the long-term establishment of neopolyploid populations. Future work that compares the overwintering ability and fate of turions between diploids and neopolyploids across multiple seasons will reveal if neopolyploids investing more into future growth in one season is advantageous in the long term. Furthermore, while our study in an asexually reproducing species can not address the roles of mating system, the current results confirm that dynamics based on asexual reproduction alone can be critical to coexistence (Rausch & Morgan 2005).

The consequences of neopolyploidy on competition at the population-level transcends a simple genome doubling effect (Doyle & Coate, 2019). Competitive interaction strength between neopolyploids and their diploid progenitors strongly depended on their genotype of origin (Fig. 4), indicating that the effect of neopolyploidy on competitive dynamics carries with it a genetic or epigenetic feature that differs among our four genetic backgrounds of neopolyploids. Because each of the four synthetic neopolyploid genetic lineages represents an independent evolutionary experiment (Bomblies, 2020), this variation suggests that genetic chance events can dictate the outcome of future inter-annual competitive dynamics and be predictive of long-term establishment of neopolyploid populations. Therefore, if genetic lineages of neopolyploid populations that produce more dormant propagules can overcome the short-term numerical costs in current growth, then establishment in may be favored in environments with a high degree of seasonality where neopolyploid populations would benefit from their greater future growth investment.

## Data availability

All data associated with this work will be freely available on the data repository Zenodo (DOI: 10.5281/zenodo.6773513).

## Acknowledgements

We thank Jae Kerstetter for assistance in maintaining the *S. polyrhiza* founding genotypes; Kayla Downs and Jason Simmons for polyploid-diploid stock creation and maintenance; Laurie Follweiler for greenhouse assistance; Ashman and Turcotte labs for feedback, discussion, and helping with experiments. This project was supported by the Dietrich

School of Arts & Sciences and grants from the National Science Foundation (#2109452 to T.J.A., #2027604 & 1912180 to T-L.A., and #1935410 to M.M.T.).

## Conflict of Interest

The authors have no conflicts of interest to report.

## Author contributions

T.J.A, T-L.A., and M.M.T. conceptualized and designed the study. E.M.O. synthesized the neopolyploids, confirmed ploidy by flow cytometry an maintained all materials. T.J.A. carried out the experiments. T.J.A. analyzed the data and wrote the first draft of the manuscript, and T.J.A., T-L.A., and M.M.T. edited subsequent drafts.

## Notes

https://zenodo.org/record/6773513#.Y2AjnOzMLlw

### Competing Interest Statement

The authors have declared no competing interest.

