## Supplemental Information for "Polyploidy impacts population growth and competition with diploids: multigenerational experiments reveal key life history tradeoffs"

**This file includes:**

Extended Methods
Figures S1 to S4
Tables S1 to S13
SI References

### Extended Methods
#### Creation and confirmation of synthetic neopolyploids

We induced whole genome duplication in the four genetically distinct diploid clones by application of colchicine (Sigma-Aldrich). to developing fronds. Briefly, we supplied independent monocultures of each diploid lineage with 1% colchicine in the presence of 0.5% DMSO and 0.03% Triton X-100 dissolved in Hoagland's growth media (Vunsh *et al.*, 2015). We also included a solvent-only control for each of the four diploid lineages by treating them with 0.5% DMSO and 0.03% Triton X-100 but did not receive 1% colchicine. Immediately following either colchicine or solvent-only control media to diploids, we placed them in a growth chamber maintained at 23˚C for 72 hours in complete darkness to avoid light-induced degradation of colchicine. Next, the colchicine media was washed from the plants with deionized water and placed in 50 % Hoagland's media until they formed populations large enough for flow cytometry to assess ploidy following the protocol in (Wei *et al.*, 2020). Specifically, we collected nuclei by chopping 20 mg of fresh frond tissue in the presence of 1 mL of Galbraith buffer solution (Galbraith *et al.*, 1983) on ice. After filtering this solution through a 40 µm pore size nylon mesh and adding 10 µL of RNase A to clean the sample of RNA, we stained the nuclei suspended in the buffer solution with a 0.1 : 1 (w/v) propidium iodide : deionized water solution. After gently mixing the propidium iodide solution and nuclei to homogenize the sample, we confirmed the ploidy level of each sample on a BD Accuri C6 cytometer by comparing the relative intensity of the peaks in relation to a diploid control sample on the FL2 channel, also requiring that the peak intensity on the FL2 channel reach a minimum of 200 events with a coefficient of variation ≤5%. We thus confirmed the neotetraploid cytotype for all four of the neotetraploid lineages used in the present study by carrying out this flow cytometry protocol twice on each neotetraploid line.

We tested for an effect of colchicine by comparing the population growth of diploids treated with colchicine to solvent-only treated diploids. In a generalized mixed effect model with a negative binomial distribution of the population sizes of these treated and untreated diploids over time, explained by colchicine, nutrients, lineage and a covariate of the initial surface area of fronds at the beginning of the experiment, there was no main effect of colchicine application ($\chi2 = 0.06$, df = 1, P = 0.940). However, there was a significant interaction between genetic lineage and colchicine application ($\chi2 = 69.56$, df = 3, P < 0.001), but there was no clear pattern in how these four diploid lineages responded to colchicine (Fig. S1). We thus excluded diploids that were not treated with colchicine from further analyses, such that we only report on the comparison between neopolyploids (generated via colchicine induction) and diploids that were treated with colchicine but failed to become polyploid. This is an important distinction since we are thus controlling for colchicine treatment by only comparing two cytotypes (diploid and neopolyploid) that have both had a history of colchicine treatment.

**Nutrient concentration and stoichiometry supply experiment**

Prior to the experiment duckweeds were pre-treated in a common garden, with quarter-strength modified Appenroth media in a growth chamber set to a 16:8 L:D cycle, maintained at constant 25˚C temperature with 50% relative humidity for three weeks, and followed standard protocols for duckweed rearing (Hart *et al.*, 2019; Subramanian & Turcotte, 2020). To investigate the effect of neopolyploidy on population performance across different resource supplies, we conducted a monoculture experiment that manipulated the supply of both nitrogen (N) and phosphorus (P) while holding all other nutrients constant. The concentration of P in the intermediate supply treatment level was based on (Appenroth *et al.*, 1996), and the corresponding supply of N at this level scaled with P so that the ratio of N : P had a mass ratio of 14, which is considered optimally colimiting for plant growth (Gusewell, 2004). We either increased or decreased the concentration of N and / or P by one order of magnitude, respectively. Thus, we had nine nutrient treatment levels, which varied in both the concentration and ratio of N and P (Table S5 for recipe). We grew 540 monoculture populations of duckweeds in a randomized block design in the greenhouse with 5 spatial blocks and one replicate per block. Of the 540 total samples, they were evenly distributed across the three ploidy levels (diploids with a history of colchicine, neopolyploids with a history of colchicine, and solvent-only diploid controls), nine nutrient levels, and four genetic lineage backgrounds, resulting in five replicates per treatment level. This experiment was conducted in a greenhouse at the University of Pittsburgh in May 2021 for 17 days under natural light conditions and a max 23ºC daytime temperature and a minimum 16ºC nighttime temperature with 35% relative humidity. To account for evaporative loss, we added 250 ml. of sterile deionized water to each container on the seventh day of the experiment.

At the start of the experiment in May 2021, we transferred six healthy fronds of each ploidy and genetic lineage into respective experimental containers using flame sterilization technique with forceps. We then photographed each container with a Nikon D3500 dSLR camera from above and at a perpendicular angle to the water surface. We included a small circular plastic size standard with a diameter of 5.5 millimeters to measure the surface area covered by the six starting fronds in the experiment. We subsequently photographed each sample unit every two days. From these photographs over the days of the experiment, we counted the number of living fronds and turions produced. At the end of the experiment on day 17, we collected all biomass from each container into a labeled and pre-weighed aluminum foil square and dried that tissue in a 55˚C drying oven for four days. To estimate the biomass investment current growth as adult fronds separately from storage as turions, we physically separated turions from actively growing fronds with forceps and placed the turions in aluminum foil squares that were distinct from the fronds for all samples in the first two blocks of the experiment.

**Competition experiment**

We followed the same pre-treatment common garden condition outlined above, prior to the start of this experiment. To test whether variation in nutrient supply can mediate the outcomes of neopolyploid competition with their diploid progenitors, we conducted an experiment where we grew diploids and neopolyploids either alone or together, in any of three possible nutrient environments in a second greenhouse experiment starting in June 2021. Since we did not detect an interaction between ploidy and nutrient stoichiometry in our first experiment, we used the three nutrient treatments with a fixed N: P stoichiometry of 14. These nutrient treatments corresponded to the lowest (N = 0.4 mg/L; P = 0.02857 mg/L), medium (N = 4 mg/L; P = 0.2857 mg/L), and highest (N = 40 mg/L; P = 2.857 mg/L) concentration of N and P in our first experiment, and we refer to these treatments in this experiment as low, medium, and high. We set this experiment up the same as described above, with the exception that for competition we followed a simple additive design (Gibson *et al.*, 1999), in which we placed six fronds of both neopolyploids and diploids with a shared genetic lineage background in the same container.

In this experiment, we had 360 experimental mesocosms which were 1 liter plastic soup containers (Webstaurant). Of the 360 sample units, they were evenly distributed across two ploidy levels, three nutrient levels, with or without competition, and four genetic lineage backgrounds, for a total of 10 replicates per treatment level. Since we did not detect a pattern of discernable differences between our solvent-only diploid control and colchicine-treated diploids, we omitted the solvent-only controls from this competition experiment. To track the provenance of diploid vs. neopolyploid daughter fronds in the containers where both ploidy levels were grown together, we applied either a small blue or red acrylic paint (Target Brand) dot with the tip of a toothpick to each respective ploidy level of daughter frond in the experiment. To maintain a balanced design, we applied a small acrylic paint dot to every plant in the experiment, regardless of being grown as a monoculture or under competition. To avoid any possible differences in the effect of blue versus red paint, we applied a red dot to half of the replicates for each treatment level and a blue dot to the remaining half of those replicates. Every other day over the experiment, we painted new daughter fronds that were still attached to their painted mothers and then photographed each experimental container.

We concluded the experiment after 17 days in the greenhouse. For samples in which there was competition and turions produced, we were unable to assign ploidy to those turions that were at the bottom of the containers and thus determined their ploidy via flow cytometry. We collected every turion from the bottom of the containers that had both diploids and neopolyploids growing and placed one turion per cell culture tray well filled with half strength Appenroth media, with a total of 299 collected turions. After one month, once the turions had germinated and produced enough biomass, we performed flow cytometry following the methods outlined above with the exception that we used 1:1 fresh trout blood: Alsever's Solution (Sigma-Aldrich) as our control standard (rather than a diploid control) and recorded their ploidy level. To account for the removal of this biomass from the experimental units on day of harvest, we estimated the amount of removed dry biomass by estimating the dry biomass per turion for each genetic lineage background and ploidy level from the first experiment with a linear regression equation and multiplying that number by the number of removed turions per ploidy. For all other samples that either did not produce turions and / or were monocultures, we measured dry biomass for each sample as in the previous experiment.

**Measurement of population demographic data**

We quantified three different measures of population performance. From the photographs, we measured population densities by counting the number of actively growing fronds over time, we similarly counted turions produced over time. We also measured productivity as dried biomass from day of harvest. To account for potential size differences between genetic lineage backgrounds and ploidy levels before treatments were applied, we measured the surface area covered by the initial six fronds in each sample from the photographs on the first day of the experiment as a covariate in all subsequent statistical models. These surface areas (in $mm^2$) were measured using ImageJ software (Schneider *et al.*, 2012).

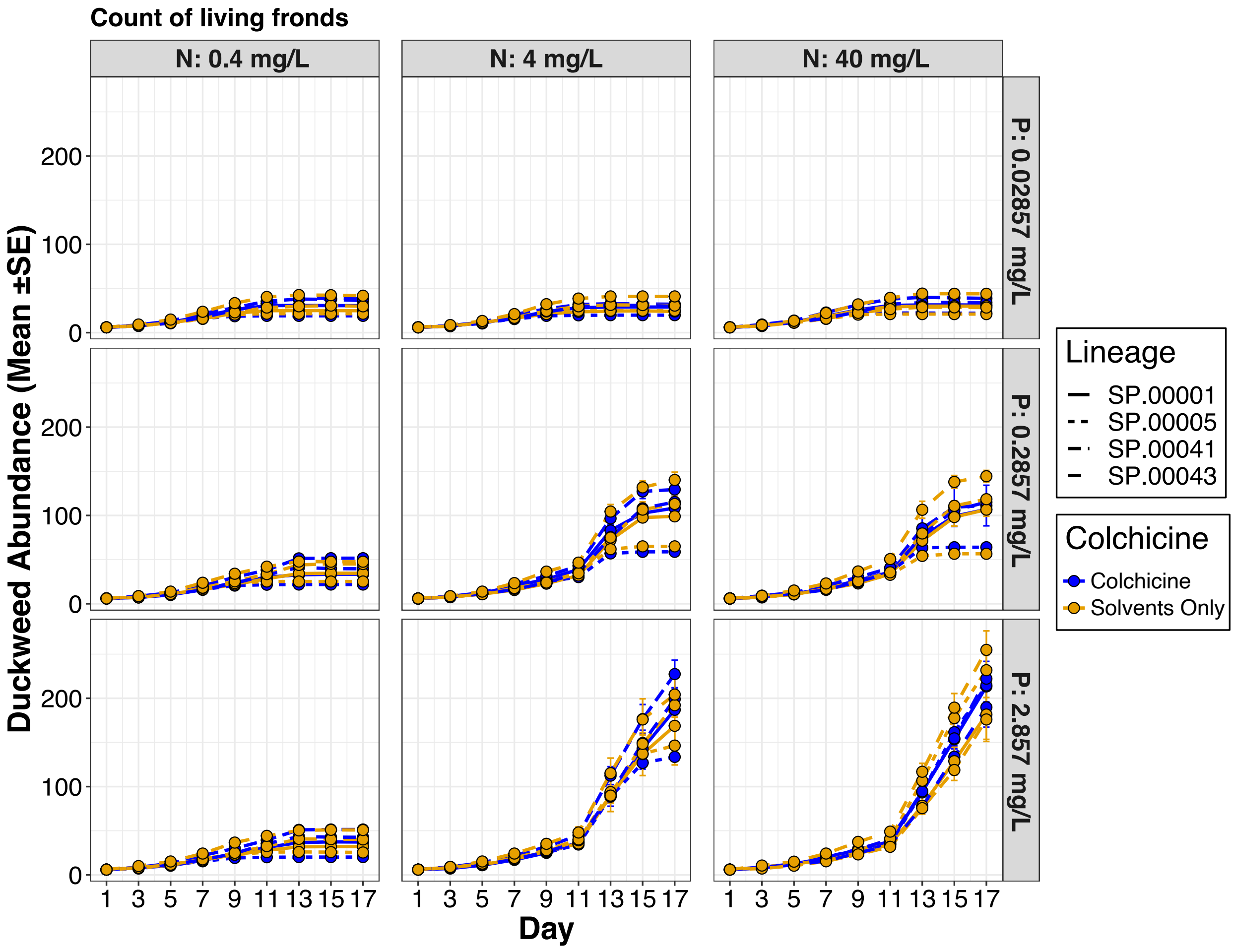


Fig. S1: Population sizes of diploids in response to either colchicine application history or treatment with solvents only (vehicle for colchicine application) over the course of the monoculture experiment. The four line types represent the four distinct genetic lineages of diploids used in this study.

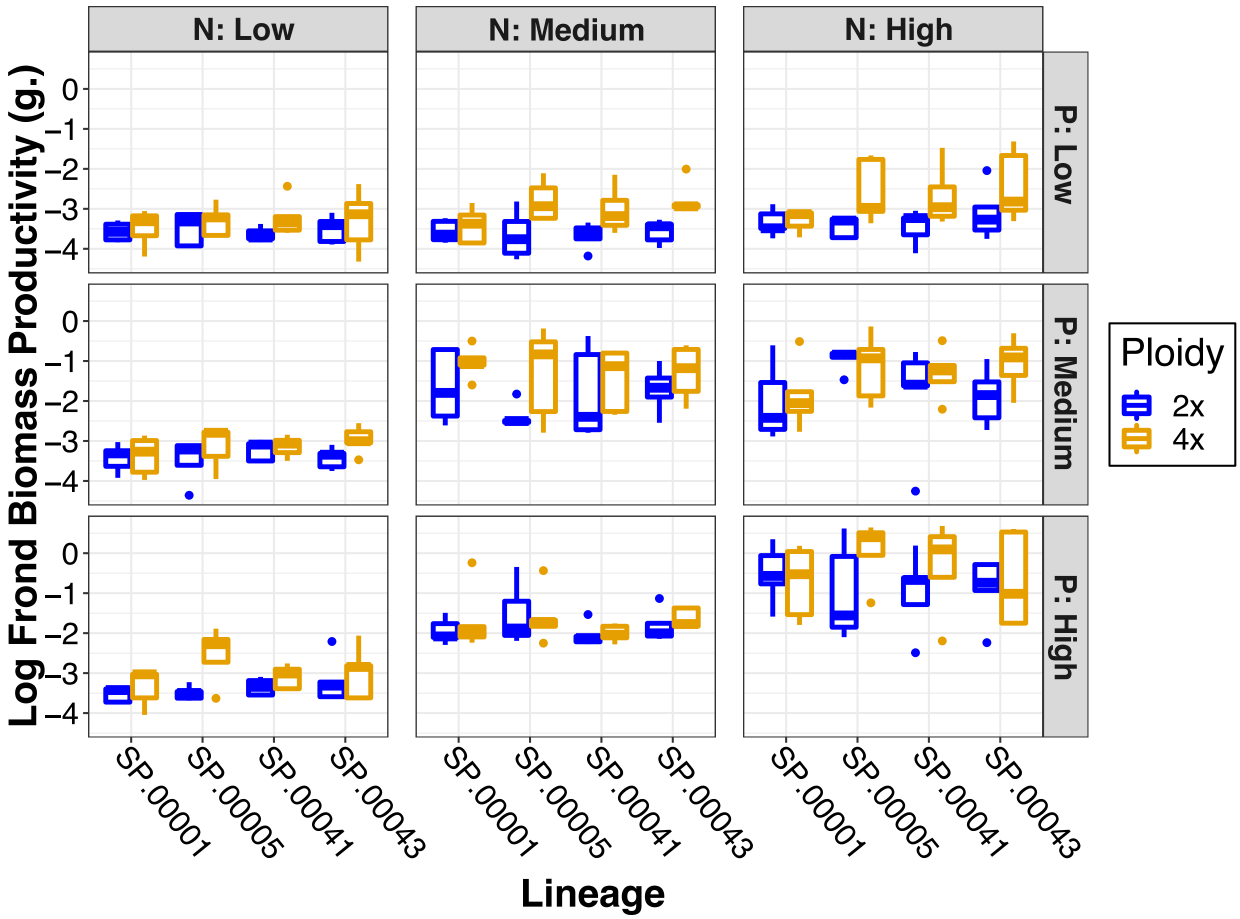


Fig. S2: Dry frond biomass productivity ($\log_{10}$) responses of the two ploidy levels for each of the four genetic lineages across the nine nutrient treatments.

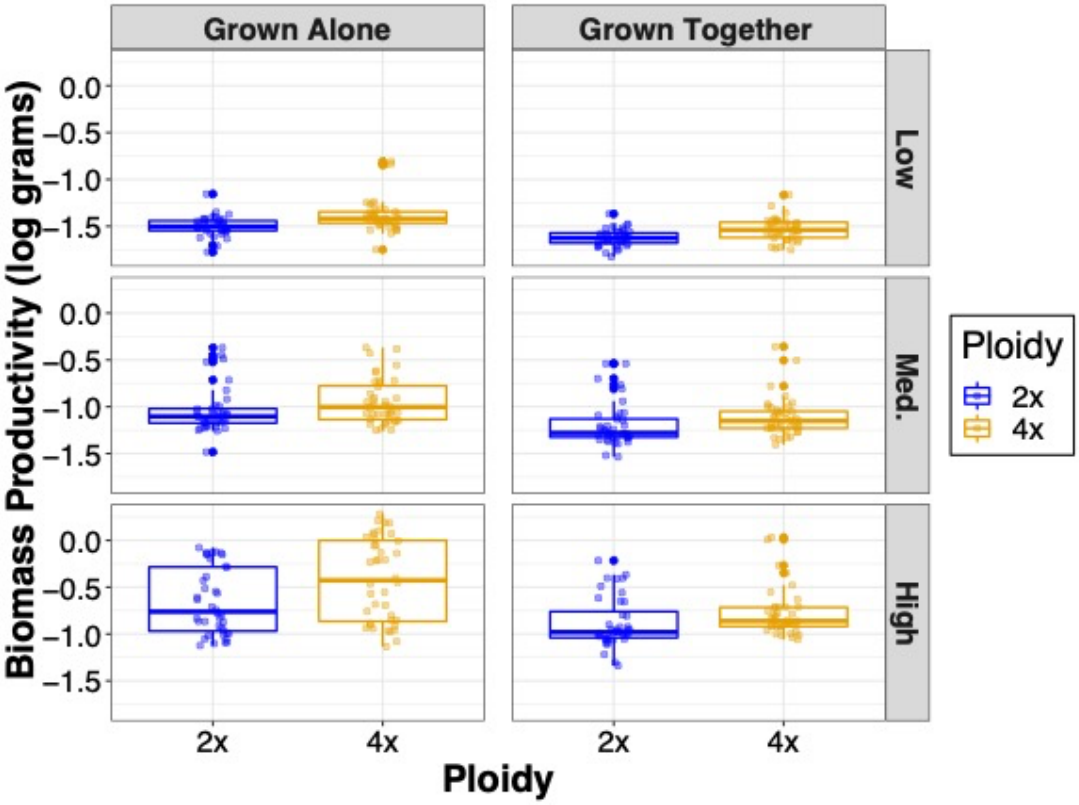


Fig. S3: Population productivity of $log_{10}$ frond dry biomass from day of harvest for diploids and neopolyploids in response to competition and nutrient enrichment. Population productivity is greater for neopolyploids than diploids, and both diploid and neopolyploid productivity are negatively affected by competition.

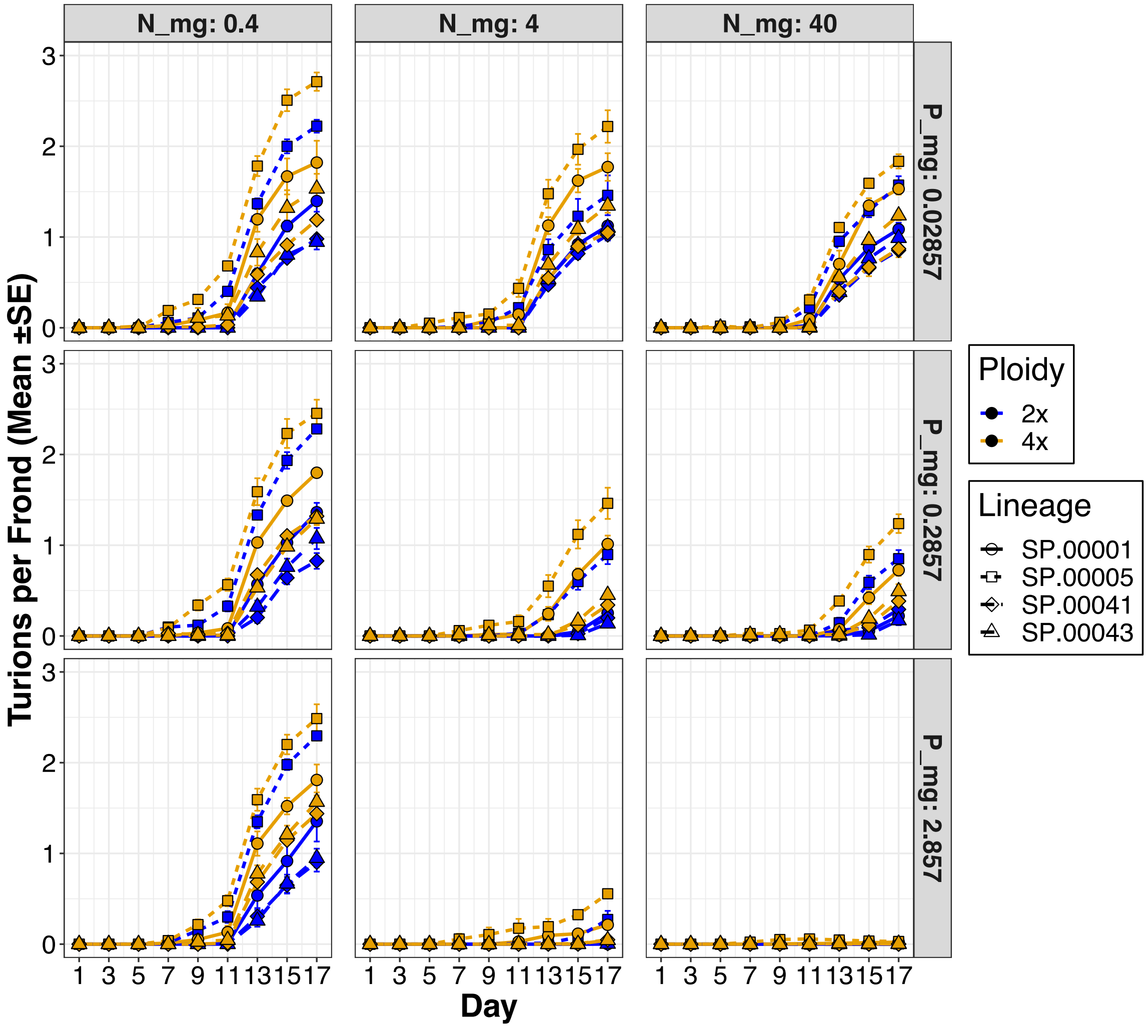


Fig. S4: Turions produced per frond in populations of diploids or neopolyploids. The four different line types with different shapes represent the four genetic lineages.

Table S1: Genetic lineages of *Spirodela polyrhiza* used in this study and their corresponding latitude and longitude of the collection site.

| Genetic Lineage Name | Latitude | Longitude |
|---|---|---|
| SP.00001 | 40.73078333 | 80.35900000 |
| SP.00005 | 40.62100000 | 79.82675000 |
| SP.00041 | 40.94098333 | 80.09125000 |
| SP.00043 | 41.60588333 | 80.53876667 |

Table S2: Nitrogen and Phosphorus treatment recipe including micronutrients in Appenroth media. The three treatment levels with N:P of 14 correspond to the low, medium, and high nutrient treatments in the competition experiment (from lowest to highest N & P concentrations).

| Nutrient Treatment | KNO3 (mg/L) | KH2PO4 (mg/L) | Nitrogen (mg/L) | Phosphorus (mg/L) | N:P |
|---:|---:|---:|---:|---:|---:|
| Low P; Low N | 2.88866 | 0.12552 | 0.4 | 0.02857 | 14 |
| Low P; Med. N | 28.88663 | 0.12552 | 4 | 0.02857 | 140 |
| Low P; High N | 288.86629 | 0.12552 | 40 | 0.02857 | 1400 |
| Med. P; Low N | 2.88866 | 1.25525 | 0.4 | 0.2857 | 1.4 |
| Med. P; Med. N | 28.88663 | 1.25525 | 4 | 0.2857 | 14 |
| Med. P; High N | 288.86629 | 1.25525 | 40 | 0.2857 | 140 |
| High P; Low N | 2.88866 | 12.55249 | 0.4 | 2.857 | 0.14 |
| High P; Med. N | 28.88663 | 12.55249 | 4 | 2.857 | 1.4 |
| High P; High N | 288.86629 | 12.55249 | 40 | 2.857 | 14 |

**Micronutrients added to all treatments**

| Reagent Name | Ionic Salt | Molar mass (g) | Concentration Stock (M) | Diluted sample concentration (mM) |
|---:|---:|---:|---:|---:|
| Potassium sulfate | K2SO4 | 174.259 | 0.040890858 | 1.022 |
| Magnesium sulfate | MgSO4-7H20 | 246.47 | 0.02 | 0.500 |
| Calcium chloride | CaCl2 | 110.98 | 0.00103 | 0.026 |
| Iron chloride | FeCL3 | 162.21 | 0.0005 | 0.013 |
| EDTA | EDTA | 292.24 | 0.0005 | 0.013 |
| Boric acid | H3BO3 | 61.83 | 0.0001 | 0.003 |
| Sodium Molybdate | Na2MoO4-2H20 | 241.95 | 0.000008 | 0.0002 |
| Manganous chloride | MnCl2-4H20 | 197.91 | 0.00026 | 0.007 |

Table S3: ANOVA table of population sizes over the days of the monoculture experiment. Significant factors are bolded with their P value.

| Factor | NumDF | DenDF | F value | P value |
|---|---|---|---|---|
| **Ploidy** | 1 | 73.23 | 12.9963 | **0.0006** |
| **Lineage** | 3 | 49.24 | 10.2034 | **<0.0001** |
| **Nitrogen** | 2 | 3035.01 | 156.257 | **<0.0001** |
| **Phosphorus** | 2 | 3035 | 241.397 | **<0.0001** |
| **Day** | 1 | 7 | 274.596 | **<0.0001** |
| **Initial Surface Area** | 1 | 3038.59 | 155.611 | **<0.0001** |
| Ploidy: Lineage | 3 | 49.04 | 1.8754 | 0.1460 |
| Ploidy: Nitrogen | 2 | 3035 | 1.6394 | 0.1943 |
| Lineage: Nitrogen | 6 | 3035 | 0.5779 | 0.7482 |
| Ploidy: Phosphorus | 2 | 3035 | 1.4667 | 0.2308 |
| Lineage: Phosphorus | 6 | 3035 | 1.1258 | 0.3444 |
| **Nitrogen: Phosphorus** | 4 | 3035 | 48.7173 | **<0.0001** |
| **Ploidy: Day** | 1 | 49 | 135.3175 | **<0.0001** |
| **Lineage: Day** | 3 | 49 | 64.9854 | **<0.0001** |
| **Nitrogen: Day** | 2 | 3035 | 1010.8274 | **<0.0001** |
| **Phosphorus: Day** | 2 | 3035 | 1406.6776 | **<0.0001** |
| Ploidy: Lineage: Nitrogen | 6 | 3035 | 0.7828 | 0.5833 |
| Ploidy: Lineage: Phosphorus | 6 | 3035 | 0.5031 | 0.8065 |
| Ploidy: Nitrogen: Phosphorus | 4 | 3035 | 0.6866 | 0.6012 |
| Lineage: Nitrogen: Phosphorus | 12 | 3035 | 0.6275 | 0.8205 |
| **Ploidy: Lineage: Day** | 3 | 49 | 3.1034 | **0.0350** |
| **Ploidy: Nitrogen: Day** | 2 | 3035 | 8.8665 | **0.0001** |
| **Lineage: Nitrogen: Day** | 6 | 3035 | 6.1683 | **<0.0001** |
| **Ploidy: Phosphorus: Day** | 2 | 3035 | 3.2742 | **0.0380** |
| **Lineage: Phosphorus: Day** | 6 | 3035 | 4.5228 | **0.0001** |
| **Nitrogen: Phosphorus: Day** | 4 | 3035 | 265.0665 | **<0.0001** |
| Ploidy: Lineage: Nitrogen: Phosphorus | 12 | 3035 | 0.4758 | 0.9298 |
| **Ploidy: Lineage: Nitrogen: Day** | 6 | 3035 | 3.0023 | **0.0063** |
| Ploidy: Lineage: Phosphorus: Day | 6 | 3035 | 1.2137 | 0.2959 |
| Ploidy: Nitrogen: Phosphorus: Day | 4 | 3035 | 2.1956 | 0.0670 |
| **Lineage: Nitrogen: Phosphorus: Day** | 12 | 3035 | 2.0491 | **0.0172** |
| Ploidy: Lineage: Nitrogen: Phosphorus: Day | 12 | 3035 | 1.4918 | 0.1194 |

Table S4: ANOVA table of estimated carrying capacities in response to polyploidy, nutrient treatment, genetic lineage, and their interactions in the monoculture experiment. Significant factors are bolded with their P value.

| **Factor** | **DF** | **F value** | **Pr(>F)** |
|---|---|---|---|
| **Ploidy** | 1 | 128.00 | **<0.0001** |
| **Nutrients** | 8 | 479.85 | **<0.0001** |
| **Lineage** | 3 | 56.56 | **<0.0001** |
| Ploidy: Nutrients | 8 | 1.34 | 0.225 |
| **Ploidy: Lineage** | 3 | 8.12 | **<0.0001** |
| Nutrients: Lineage | 24 | 1.17 | 0.2683 |
| Ploidy: Nutrients: Lineage | 24 | 1.15 | 0.2846 |
| Residuals | 288 | | |

Table S5: ANOVA table of population sizes over the days of the monoculture experiment. Significant factors are bolded with their P value.

| Factor | NumDF | DenDF | F value | P value |
|---|---|---|---|---|
| **Ploidy** | 1 | 3199.9 | 4.4181 | **0.0356** |
| **Lineage** | 3 | 3196.1 | 5.2449 | **0.0013** |
| **NP_ratio** | 1 | 3196 | 27.5751 | **<0.0001** |
| **Day** | 1 | 7 | 274.5962 | **<0.0001** |
| **Initial Surface Area** | 1 | 3187 | 34.1854 | **<0.0001** |
| Ploidy: Lineage | 3 | 3196 | 0.964 | 0.4087 |
| Ploidy: NP | 1 | 3196 | 0.3788 | 0.5383 |
| Lineage: NP _ratio | 3 | 3196 | 0.0262 | 0.9943 |
| **Ploidy: Day** | 1 | 3196 | 69.6346 | **<0.0001** |
| **Lineage: Day** | 3 | 3196 | 33.4416 | **<0.0001** |
| **NP_ratio: Day** | 1 | 3196 | 137.1712 | **<0.0001** |
| Ploidy: Lineage: NP_ratio | 3 | 3196 | 0.0953 | 0.9626 |
| Ploidy: Lineage: Day | 3 | 3196 | 1.597 | 0.1880 |
| Ploidy: NP_ratio: Day | 1 | 3196 | 1.2589 | 0.2619 |
| Lineage: NP_ratio: Day | 3 | 3196 | 0.0559 | 0.9826 |
| Ploidy: Lineage: NP_ratio: Day | 3 | 3196 | 0.0905 | 0.9653 |

Table S6: Chi square table of a generalized linear mixed model with a gamma distribution of dry biomass productivity in the monoculture experiment. Significant factors are bolded with their P value.

| **Factor** | **Chisq** | **Df** | **Pr(>Chisq)** |
|---|---|---|---|
| **Ploidy** | 6.2297 | 1 | **0.0126** |
| **Nitrogen** | 76.3034 | 1 | **<0.0001** |
| **Phosphorus** | 34.4402 | 1 | **<0.0001** |
| Lineage | 1.1722 | 3 | 0.7597 |
| Initial Surface Area | 0.8522 | 1 | 0.3559 |
| Ploidy: Nitrogen | 0.0284 | 1 | 0.8661 |
| Ploidy: Phosphorus | 0.2835 | 1 | 0.5944 |
| **Nitrogen: Phosphorus** | 21.2234 | 1 | **<0.0001** |
| Ploidy: Lineage | 1.6222 | 3 | 0.6544 |
| Nitrogen: Lineage | 2.8824 | 3 | 0.4101 |
| Phosphorus: Lineage | 1.4964 | 3 | 0.6831 |
| Ploidy: Nitrogen: Phosphorus | 0.2645 | 1 | 0.6070 |
| Ploidy: Nitrogen: Lineage | 2.2825 | 3 | 0.5159 |
| Ploidy: Phosphorus: Lineage | 0.8287 | 3 | 0.8426 |
| Nitrogen: Phosphorus: Lineage | 2.1211 | 3 | 0.5477 |
| Ploidy: Nitrogen: Phosphorus: Lineage | 1.0951 | 3 | 0.7783 |

Table S7: ANOVA table of population sizes over the days of the competition experiment. Significant factors are bolded with their P value.

| **Factor** | **NumDF** | **DenDF** | **F value** | **Pr(>F)** |
|---|---:|---:|---:|---:|
| Ploidy | 1 | 35 | 0.061 | 0.8057 |
| **Nutrients** | 2 | 3224 | 488.578 | **<0.0001** |
| **Competition** | 1 | 3224 | 16.560 | **<0.0001** |
| **Day** | 1 | 5 | 289.779 | **<0.0001** |
| **Lineage** | 3 | 35 | 5.963 | **0.0021** |
| Ploidy: Nutrients | 2 | 3224 | 0.144 | 0.8661 |
| Ploidy: Competition | 1 | 3224 | 0.590 | 0.4424 |
| **Nutrients: Competition** | 2 | 3224 | 12.588 | **<0.0001** |
| **Ploidy: Day** | 1 | 35 | 192.887 | **<0.0001** |
| **Nutrients: Day** | 2 | 3224 | 2780.591 | **<0.0001** |
| **Competition: Day** | 1 | 3224 | 80.521 | **<0.0001** |
| Ploidy: Lineage | 3 | 35 | 2.152 | 0.1112 |
| Nutrients: Lineage | 6 | 3224 | 1.341 | 0.2353 |
| Competition: Lineage | 3 | 3224 | 0.139 | 0.9365 |
| **Day: Lineage** | 3 | 35 | 15.216 | **<0.0001** |
| Ploidy: Nutrients: Competition | 2 | 3224 | 0.405 | 0.6672 |
| **Ploidy: Nutrients: Day** | 2 | 3224 | 8.437 | **0.0002** |
| **Ploidy: Competition: Day** | 1 | 3224 | 6.920 | **0.0086** |
| **Nutrients: Competition: Day** | 2 | 3224 | 48.050 | **<0.0001** |
| Ploidy: Nutrients: Lineage | 6 | 3224 | 0.291 | 0.9415 |
| Ploidy: Competition: Lineage | 3 | 3224 | 0.261 | 0.8536 |
| Nutrients: Competition: Lineage | 6 | 3224 | 0.402 | 0.8783 |
| Ploidy: Day: Lineage | 3 | 35 | 1.909 | 0.1462 |
| **Nutrients: Day: Lineage** | 6 | 3224 | 4.770 | **<0.0001** |
| Competition: Day: Lineage | 3 | 3224 | 1.276 | 0.2809 |
| Ploidy: Nutrients: Competition: Day | 2 | 3224 | 0.322 | 0.7249 |
| Ploidy: Nutrients: Competition: Lineage | 6 | 3224 | 0.342 | 0.9148 |
| Ploidy: Nutrients: Day: Lineage | 6 | 3224 | 1.017 | 0.4118 |
| **Ploidy: Competition: Day: Lineage** | 3 | 3224 | 2.877 | **0.0348** |
| Nutrients: Competition: Day: Lineage | 6 | 3224 | 0.689 | 0.6589 |
| Ploidy: Nutrients: Competition: Day: Lineage | 6 | 3224 | 0.614 | 0.7192 |

Table S8: Chi square table of a general linear estimated carrying capacities in the competition experiment. This analysis omits samples grown in high nutrients because their population sizes did not reach an asymptote and did not fit the logistic model. Significant factors are bolded with their P value.

| Factor | Chisq | Df | Pr(>Chisq) |
|---|---|---|---|
| **Ploidy** | 7.98 | 1 | **0.0047** |
| **Nutrients** | 366.48 | 1 | **0.0000** |
| **Competition** | 6.59 | 1 | **0.0103** |
| Lineage | 3.49 | 3 | 0.3221 |
| Ploidy: Nutrients | 0.13 | 1 | 0.7171 |
| Ploidy: Competition | 0.28 | 1 | 0.5968 |
| Nutrients: Competition | 2.02 | 1 | 0.1550 |
| Ploidy: Lineage | 1.11 | 3 | 0.7740 |
| Nutrients: Lineage | 4.93 | 3 | 0.1772 |
| Competition: Lineage | 1.75 | 3 | 0.6258 |
| Ploidy: Nutrients: Competition | 0.28 | 1 | 0.5965 |
| Ploidy: Nutrients: Lineage | 1.21 | 3 | 0.7507 |
| Ploidy: Competition: Lineage | 0.48 | 3 | 0.9223 |
| Nutrients: Competition: Lineage | 0.79 | 3 | 0.8518 |
| Ploidy: Nutrients: Competition: Lineage | 1.34 | 3 | 0.7206 |

Table S9: *Post-hoc* least square means estimates of biomass productivity from the harvested tissues in the competition experiment. The ± 95 % confidence interval is indicated by the asymp.LCI and asymp.UCI columns, respectively.

| Ploidy | Nutrients | Competition | LS Mean | SE | asymp.LCI | asymp.UCI |
|---|---|---|---|---|---|---|
| 2x | High | Monoculture | 0.314 | 0.02961 | 0.256 | 0.3721 |
| 4x | High | Monoculture | 0.5888 | 0.05661 | 0.4779 | 0.6998 |
| 2x | Low | Monoculture | 0.0387 | 0.00382 | 0.0312 | 0.0462 |
| 4x | Low | Monoculture | 0.0415 | 0.0047 | 0.0323 | 0.0507 |
| 2x | Med. | Monoculture | 0.1126 | 0.01048 | 0.0921 | 0.1332 |
| 4x | Med. | Monoculture | 0.1321 | 0.01321 | 0.1062 | 0.158 |
| 2x | High | Competition | 0.1678 | 0.01612 | 0.1362 | 0.1993 |
| 4x | High | Competition | 0.2039 | 0.02294 | 0.1589 | 0.2488 |
| 2x | Low | Competition | 0.0299 | 0.00308 | 0.0238 | 0.0359 |
| 4x | Low | Competition | 0.0233 | 0.0034 | 0.0166 | 0.0299 |
| 2x | Med. | Competition | 0.0815 | 0.00781 | 0.0662 | 0.0968 |
| 4x | Med. | Competition | 0.0848 | 0.00918 | 0.0669 | 0.1028 |

Table S10: Chi square table of a generalized linear mixed model of biomass productivity in the competition experiment with a gamma distribution. Significant factors are bolded with their P value.

| **Factor** | **Chisq** | **Df** | **Pr(>Chisq)** |
|---|---|---|---|
| Ploidy | 0.1655 | 1 | 0.6842 |
| **Nutrients** | 370.0922 | 2 | **<0.0001** |
| **Competition** | 29.5635 | 1 | **<0.0001** |
| Lineage | 1.5101 | 3 | 0.6800 |
| **Initial Surface Area** | 8.785 | 1 | **0.0030** |
| Ploidy: Nutrients | 3.3601 | 2 | 0.1864 |
| **Ploidy: Competition** | 3.9774 | 1 | **0.0461** |
| **Nutrients: Competition** | 49.7983 | 2 | **<0.0001** |
| Ploidy: Lineage | 0.4232 | 3 | 0.9354 |
| **Nutrients: Lineage** | 12.9847 | 6 | **0.0433** |
| Competition: Lineage | 1.545 | 3 | 0.6719 |
| **Ploidy: Nutrients: Competition** | 13.4171 | 2 | **0.0012** |
| Ploidy: Nutrients: Lineage | 8.4003 | 6 | 0.2102 |
| Ploidy: Competition: Lineage | 1.5112 | 3 | 0.6797 |
| Nutrients: Competition: Lineage | 11.1496 | 6 | 0.0839 |
| Ploidy: Nutrients: Competition: Lineage | 7.1068 | 6 | 0.3111 |

Table S11: Chi square table of a generalized linear mixed model with binomial distribution of the timing of turion production (day that turions were present or absent) in the monoculture experiment. Significant factors are bolded with their P value.

| **Factor** | **Chisq** | **Df** | **Pr(>Chisq)** |
|---|---:|---:|---:|
| **Ploidy** | 25.8695 | 7 | **0.0005** |
| **Lineage** | 141.7313 | 15 | **<0.0001** |
| **N** | 65.2875 | 7 | **<0.0001** |
| **P** | 75.059 | 6 | **<0.0001** |
| **Day** | 203.436 | 7 | **<0.0001** |
| **Initial Surface Area** | 6.2769 | 1 | **0.0122** |
| **Ploidy: Lineage** | 20.7892 | 8 | **0.0077** |
| **Ploidy: N** | 10.6821 | 4 | **0.0304** |
| **Lineage: N** | 15.8782 | 8 | **0.0442** |
| **Ploidy: P** | 11.6082 | 3 | **0.0089** |
| Lineage: P | 11.5639 | 7 | 0.1158 |
| **N: P** | 9.5413 | 3 | **0.0229** |
| **Ploidy: Day** | 8.0826 | 3 | **0.0443** |
| Lineage: Day | 11.894 | 7 | 0.1041 |
| N: Day | 5.2561 | 2 | 0.0722 |
| **P: Day** | 9.8398 | 2 | **0.0073** |
| Ploidy: Lineage: N | 8.5855 | 5 | 0.1268 |
| **Ploidy: Lineage: P** | 9.6879 | 4 | **0.0460** |
| **Ploidy: N: P** | 10.3956 | 2 | **0.0055** |
| **Lineage: N: P** | 12.6434 | 4 | **0.0132** |
| Ploidy: Lineage: Day | 5.7401 | 3 | 0.1250 |
| **Ploidy: N: Day** | 6.0076 | 2 | **0.0496** |
| Lineage: N: Day | 8.753 | 4 | 0.0676 |
| Ploidy: P: Day | 0.9396 | 1 | 0.3324 |
| Lineage: P: Day | 4.03 | 3 | 0.2582 |
| N: P: Day | 0.3742 | 1 | 0.5407 |
| **Ploidy: Lineage: N: P** | 13.1321 | 3 | **0.0044** |
| Ploidy: Lineage: N: Day | 7.1316 | 3 | 0.0678 |
| Ploidy: Lineage: P: Day | 3.5055 | 3 | 0.3201 |
| Ploidy: N: P: Day | 0.2917 | 1 | 0.5891 |
| Lineage: N: P: Day | 2.6109 | 3 | 0.4556 |
| Ploidy: Lineage: N: P: Day | 7.4422 | 3 | 0.0591 |

Table S12: ANOVA table of a linear mixed model of $log_{10}$ transformed (log +1) total turion production over the course of the monoculture experiment. Significant factors are bolded with their P value.

| Factor | NumDF | DenDF | F value | P value |
|---|---|---|---|---|
| Ploidy | 1 | 56.89 | 3.3913 | 0.0708 |
| Lineage | 3 | 49.08 | 0.3199 | 0.8109 |
| **Nitrogen** | 2 | 3035.17 | 46.5418 | **<0.0001** |
| **Phosphorus** | 2 | 3035.11 | 89.106 | **<0.0001** |
| **Day** | 1 | 7 | 29.202 | **0.0010** |
| **Initial Surface Area** | 1 | 1900.44 | 17.9942 | **<0.0001** |
| Ploidy: Lineage | 3 | 49.01 | 0.2528 | 0.8590 |
| **Ploidy: Nitrogen** | 2 | 3034.99 | 5.7792 | **0.0031** |
| **Lineage: Nitrogen** | 6 | 3035.01 | 4.6954 | **<0.0001** |
| **Ploidy: Phosphorus** | 2 | 3034.98 | 5.6625 | **0.0035** |
| **Lineage: Phosphorus** | 6 | 3035.05 | 4.0361 | **0.0005** |
| **Nitrogen: Phosphorus** | 4 | 3035 | 33.8106 | **<0.0001** |
| **Ploidy: Day** | 1 | 49 | 16.8846 | **0.0002** |
| **Lineage: Day** | 3 | 49 | 17.9907 | **<0.0001** |
| **Nitrogen: Day** | 2 | 3034.96 | 416.1398 | **<0.0001** |
| **Phosphorus: Day** | 2 | 3034.96 | 541.791 | **<0.0001** |
| Ploidy: Lineage: Nitrogen | 6 | 3035.02 | 0.9335 | 0.4695 |
| Ploidy: Lineage: Phosphorus | 6 | 3035.02 | 0.7603 | 0.6012 |
| Ploidy: Nitrogen: Phosphorus | 4 | 3035.04 | 0.7752 | 0.5412 |
| Lineage: Nitrogen: Phosphorus | 12 | 3035.04 | 1.5753 | 0.0916 |
| Ploidy: Lineage: Day | 3 | 49 | 1.4517 | 0.2392 |
| **Ploidy: Nitrogen: Day** | 2 | 3034.96 | 23.8585 | **<0.0001** |
| **Lineage: Nitrogen: Day** | 6 | 3034.96 | 8.8779 | **<0.0001** |
| **Ploidy: Phosphorus: Day** | 2 | 3034.96 | 20.5417 | **<0.0001** |
| **Lineage: Phosphorus: Day** | 6 | 3034.96 | 10.0914 | **<0.0001** |
| **Nitrogen: Phosphorus: Day** | 4 | 3034.96 | 159.7624 | **<0.0001** |
| Ploidy: Lineage: Nitrogen: Phosphorus | 12 | 3035.02 | 0.3798 | 0.9710 |
| **Ploidy: Lineage: Nitrogen: Day** | 6 | 3034.96 | 3.1336 | **0.0046** |
| **Ploidy: Lineage: Phosphorus: Day** | 6 | 3034.96 | 3.5196 | **0.0018** |
| **Ploidy: Nitrogen: Phosphorus: Day** | 4 | 3034.96 | 4.4508 | **0.0014** |
| **Lineage: Nitrogen: Phosphorus: Day** | 12 | 3034.96 | 4.9978 | **<0.0001** |
| Ploidy: Lineage: Nitrogen: Phosphorus: Day | 12 | 3034.96 | 1.5528 | 0.0985 |

Table S13: Chi square table of a generalized linear mixed model with a poisson distribution of total turion production (count) over the course of the competition experiment. Significant factors are bolded with their P value.

| Factor | Chisq | Df | Pr(>Chisq) |
|---|---|---|---|
| **Ploidy** | 298.8735 | 17 | **<0.0001** |
| **Nutrients** | 1328.0646 | 34 | **<0.0001** |
| **Day** | 531.2992 | 15 | **<0.0001** |
| **Competition** | 272.2497 | 17 | **<0.0001** |
| **Lineage** | 875.3831 | 21 | **<0.0001** |
| **Ploidy: Nutrients** | 85.6308 | 16 | **<0.0001** |
| **Ploidy: Day** | 209.5507 | 7 | **<0.0001** |
| **Nutrients: Day** | 520.9554 | 14 | **<0.0001** |
| **Ploidy: Competition** | 73.4202 | 8 | **<0.0001** |
| **Nutrients: Competition** | 118.666 | 16 | **<0.0001** |
| **Day: Competition** | 61.5676 | 7 | **<0.0001** |
| **Ploidy: Lineage** | 101.6981 | 11 | **<0.0001** |
| **Nutrients: Lineage** | 100.4687 | 21 | **<0.0001** |
| **Day: Lineage** | 141.0202 | 9 | **<0.0001** |
| **Competition: Lineage** | 60.982 | 11 | **<0.0001** |
| **Ploidy: Nutrients: Day** | 59.7074 | 6 | **<0.0001** |
| **Ploidy: Nutrients: Competition** | 58.0001 | 7 | **<0.0001** |
| **Ploidy: Day: Competition** | 70.5211 | 3 | **<0.0001** |
| **Nutrients: Day: Competition** | 54.0889 | 6 | **<0.0001** |
| **Ploidy: Nutrients: Lineage** | 33.027 | 11 | **0.0005** |
| **Ploidy: Day: Lineage** | 87.3176 | 5 | **<0.0001** |
| **Nutrients: Day: Lineage** | 39.0511 | 10 | **<0.0001** |
| **Ploidy: Competition: Lineage** | 23.6854 | 6 | **0.0006** |
| **Nutrients: Competition: Lineage** | 32.3259 | 11 | **0.0007** |
| **Day: Competition: Lineage** | 19.48 | 5 | **0.0015639** |
| **Ploidy: Nutrients: Day: Competition** | 49.3271 | 2 | **<0.0001** |
| **Ploidy: Nutrients: Day: Lineage** | 16.7754 | 6 | **0.0101** |
| Ploidy: Nutrients: Competition: Lineage | 10.1584 | 6 | 0.1181 |
| Ploidy: Day: Competition: Lineage | 2.7185 | 3 | 0.4371 |
| Nutrients: Day: Competition: Lineage | 10.6044 | 6 | 0.1014 |
| Ploidy: Nutrients: Day: Competition: Lineage | 0.0001 | 6 | 1 |
